# Characterization of a Novel Model of Overuse-Induced Calcific Achilles Tendinopathy in Mice: Contralateral Tendinopathy Following Unilateral Tenotomy

**DOI:** 10.1101/209247

**Authors:** Liang Wang, Jie Zhang, Gang-hui Yin, Zhong-min Zhang, Tian-yu Chen, Jian Jin, Ping-lin Lai, Bin Huang, Bo Yan, Yu-hui Chen, Da-di Jin, Min-jun Huang

**Affiliations:** Department of Orthopedics, the Third Affiliated Hospital of Southern Medical University, Guangzhou 510665 Guangdong, PR China; Academy of Orthopedics, Guangdong Province, Guangzhou 510665, Guangdong, PR USA; Department of Cell Biology, School of Basic Medical Science, Southern Medical University, Guangzhou 510515, Guangdong, PR China; Department of Spine Surgery, Nanfang Hospital, Southern Medical University, Guangzhou 510515, Guangdong, PR China

## Abstract

**Objective:** To develop a simple but reproducible overuse induced animal model of Achilles tendinopathy in mice for better understanding the underlying mechanism and prevention of calcific Achilles tendinopathy.

**Methods:** 80 C57/B6 mice (8-9 weeks old) were employed and randomly divided into control group and experimental group. Unilateral Achilles tenotomy was performed on the right hindlimb of experiment group. After 12 weeks, the onset of Achilles tedinopathy in the contralateral Achilles tendon was determined by radiological assessment, histological analysis, electron microscopy observation and biomechanical test.

**Results:** The onset of calcific Achilles tendinopathy in contralateral Achilles tendon was confirmed after 12 weeks unilateral tenotomy. The contralateral Achilles tendon of experimental group was characterized as hypercelluarity, neovascularization and fused collagen fiber disarrangement, compared to the control group. Importantly, intratendon endochondral ossification and calcaneus deformity was featured in contralateral Achilles tendon. Additionally, poor biomechanical properties in the contralateral Achilles tendon revealed the incidence of Achilles tedinopathy.

**Conclusion:** We hereby introduce a novel simple but reproducible spontaneous contralateral calcific Achilles tendinopathy model in mice, which represents the overuse conditions during the tendinopathy development in human-beings. It should be a useful tool to further study the underlying pathogenesis of calcific Achilles tendinopathy.

## Introduction

Achilles tendinopathy is a significant clinical disease featured by activity-related pain, focal movement limitation and intratendinous imaging changes with high prevalence in elite athletes and active population[1]. However, the treatment of Achilles tendinopathy was mainly based on theoretical rationale and clinical experience due to its unclear underlying pathogenesis[2]. Despite recent studies have recommended the Achilles tendon degeneration (tendinosis) is typical histopathological appearance during the pathogenesis of tendinopathy[3–5], its pathophysiology, the very foundation of the treatment decision, is not fully understood. On account of potential demand to further indentify the efficiency of medical intervention, validated animal models with Achilles tendinopathy should be developed to represent the consistent conditions and known risk factors in human[6]. Thus, several animal models of Achilles tendinopathy have been established in the previous studies. Sullo *et al.* administered weekly injections of PGE1 into the Achilles tendons of rats for 5 weeks to induce Achilles tendon tendinosis[7]. Moreover, Glazebrook *et al* recently employed an uphill treadmill rat model to develop Achilles tendinopathy[8]. Interestingly, Nakagawa *et al* reported a disuse model with tail suspension on rats for five weeks resulted in a decrease in collagen fiber surface area and an increase in proportion of thin to thick collagen fibers in Achilles tendon[9].

Even though existed studies shed light on the novel established models of Achilles tendinopathy, they do have distinct drawbacks such as complex inducible conditions and special treadmill instrument requirement, confounding variables interference, which may compromise its reproducible characterization and restrict its applications[10–12]. Therefore, the established animal model with more simple and high reproducible design could gain more popularity in consistent with the increasing clinical interest of Achilles tendinopathy.

In addition, since the aberrant activated differentiation of tendon-derived stem cells (TDSCs) have been supposed as a trigger of tendinopathy development, its erroneous chondrogenic and osteogenic differentiation may contribute to terminal calcific Achilles tendinopathy[13]. But the underlying mechanisms and its prevention are still poorly understood because of the absence of suitable animal model up to now. An excellent *in vivo* model will be beneficial to better understanding the underlying pathophysiology of tenocyte differentiation.

Herein, we introduce a novel Achilles tendinopathy model in mice characterized by simple procedure but highly reproducible. In this model, the disorganization of collagen fibers, neovascularity, hypercelluarity, endochondral ossification and poor Achilles tendon mechanical properties have been indentified, proven to be a successful establishment of the calcific Achilles tendinopathy.

## Materials and methods

### Animal experiment

The Animal Care and Use Committee of Southern Medical University approved all animal experimental protocols, which followed the principles expressed in the National Institute of Health Guide. Eighty male C57/B6 mice (8–9 weeks old) were obtained from the experimental animal research center of Southern Medical University. The animals were randomly divided into 2 groups (n = 40). All of the mice were anesthetized with a mixture of ketamine (90 mg/kg, i.p.), and xylazine (10 mg/kg,i. p.), and then the experimental group underwent midpoint Achilles tenotomy on left hindlimbs through a posterior approach under aseptic condition. Incision was routinely closed with an interrupted 4-0 silk suture, whilst the control group underwent skin incision only. The mice were kept in the standard condition. At 12 weeks post-operation, all animals were sacrificed by administration of a fatally high dose of anesthetics. The right Achilles tendon tissues from soleus muscle attachment to calcaneus insertional endpoint were harvested.

### Scanning electron microscopy (SEM) observation

The Achilles tendon tissues were longitudinally incised directly, and then the specimens were fixed in 2.5 % glutaraldehyde for 12 hours. They were dehydrated in a graded acetone series and critical-point-dried in carbon dioxide. After this procedure, specimens were cut into 0.5 cm × 1.0 cm pieces, fixed onto a scanning electron microscopy (SEM) stub, and sputter-coated with gold of approximately 20 nm. The images of the specimens were captured by SEM (Hitachi, S-3000N, Japan) in high vacuum mode with accelerating voltages of 10-20 kV

### Transmission electron microscopy (TEM) observation

The Achilles tendon tissue were dissected and immediately fixed in 2.5% glutaraldehyde for 1h, rinsed three times in 0.1 M sodium cacodylate buffer (pH 7.4) and re-fixed in 2% osmium oxide in 0.1 M cacodylate buffer for 2 h, then washed three times in distilled water. After that, the specimens were stained with ethanolic uranyl acetate overnight and then rinsed in distilled water, dehydrated in graded concentrations of ethanol series at 4 °C. The Achilles tendons were embedded with propylene oxide in a fresh mixture of Epon and ethanol, and polymerized for 2 days at 60 °C. Thin 50–70 nm thickness sections were cut along the longitudinal axis of the tendon with a diamond knife mounted on an ultra microtome (Meyco-Diatome, Switzerland) and placed on electron microscope grids. Sections on the grids were stained with 2% uranyl acetate and lead citrate solution and viewed in a transmission electron microscope(Hitachi, H600, Japan) operated at an accelerating voltage of 80 kV. The fibril diameters were measured, and the distribution was analyzed.

### Histological analysis

The pictures of gross examination for Achilles tendons were documented once the animals were sacrificed and limbs skinned. The Achilles tendons were isolated and fixed in 4% neutral-buffered paraformaldehyde overnight, followed by decalcified in EDTA(0.5M, PH=7.4) at 4°C for 2 weeks. Serially dehydration in graded ethanol and paraffin embedding were subsequently conducted. The blocks were cut into 4- μ m-thick sections. Hematoxylin-eosin (H&E) staining and Toluidine-blue staining were then accomplished. The celluarity and vascularity has been calculated as previously described[4,14].

For immunohistochemica((IHC) staining to evaluate the endochondral ossification in the degenerated Achilles tendons, paraffin-embedded sections were deparaffinated and rehydrated through graded ethanol. Endogenous peroxidase activity was blocked with 0.3% hydrogen peroxide in PBS. Then the sections were blocked with 5% horse serum albumin and incubated with the primary antibody of collagen II (1:100 dilution; Abcam, USA) and collagen X(1:100 dilution; Abcam, USA) at 4°C overnight. The sections were washed with PBS for three times and then incubated with secondary anti-mouse IgG for 1 h at 37°C. The colorization developed in 3, 30-diaminobenzidine tetrachloride solution.

For immunofluorescence(IF) staining, paraffin-embedded samples were deparaffinated and rehydrated in graded ethanol and water. After washing with PBS, sections were blocked with 5% horse serum for 30 min at room temperature and then incubated with mouse anti-Collagen III polyclonal antibody (1:100) (Abcam, USA) overnight at 4°C. After washing with PBS, sections were incubated with Alexa Fluor 594 goat anti mouse immunoglobulin G (ZSGB-Bio, China) for 30 min. Finally, slides were washed and mounted with prolong gold antifade reagent (Invitrogen, USA). The images were captured by confocal laser scanning microscopy (Olympus FV1000, Japan)

### Radiological analysis

All animals were assessed by X-ray analysis to detect imaging changes in Achilles tendons at 12 weeks after tenotomy. When the animals were sacrificed, the hindlimbs were amputated from knee joint for the micro-CT assessment (ZKKS-MCT-Sharp-III scanner, Caskaisheng, CHINA). The scanning system was set to 70kV, 30W, and 429μA. The 3 dimensional reconstruction of hindlimbs was conducted by 3DMed 2.2 software. The ossification or mineralization in Achilles tendons was indentified as the Region Of Interest (ROI).

### Biomechanical analysis

Tensile loading tests were performed on an Instron 5843 tensile tester (Instron, USA). The Right Achilles tendons with calcaneus were harvested at 12 weeks after tenotomy and stored in PBS at − 80°C. The cross-sectional area was assessed by measuring the thickness in three points along the tendon’s length. Then the tendons with calcaneus have been firmly fixed into clamps of the tensile tester and kept moist in PBS during the experiments. The tendons were stretched with a constant strain rate. The displacement and force has been recorded until the Achilles tendons break in the midpoint. The failure tensile load and the stiffness of Achilles tendon were then calculated.

### Statistical analysis

Data analysis was performed with SPSS 13.0. The results were presented as the mean ± SD. Two-sample t-tests were used for data analysis. P values <0.05 were considered statistically significant.

## Results

### Hypercelluarity, neovascularization in the contralateral Achilles tendon after 12 weeks unilateral tenotomy

Since hypercelluarity and neovascularization in tendon are common characters of tendinosis or tendinopathy[15], we firstly evaluated the potential changes in the contralateral Achilles tendon by H&E staining after 12 weeks unilateral tenotomy. Compared to control, there were increased and disorganized tenocytes inside the contralateral Achilles tendon. A large mount of vascular were infiltrated in the contralateral Achilles tendon (Fig.1). These results indicated that the contrallateral Achilles tendon could be hypercelluar and hypervascular after 12 weeks unilateral tenotomy, which implies the underlying tendinosis or tendinopathy incidence.

**Figure. 1.**
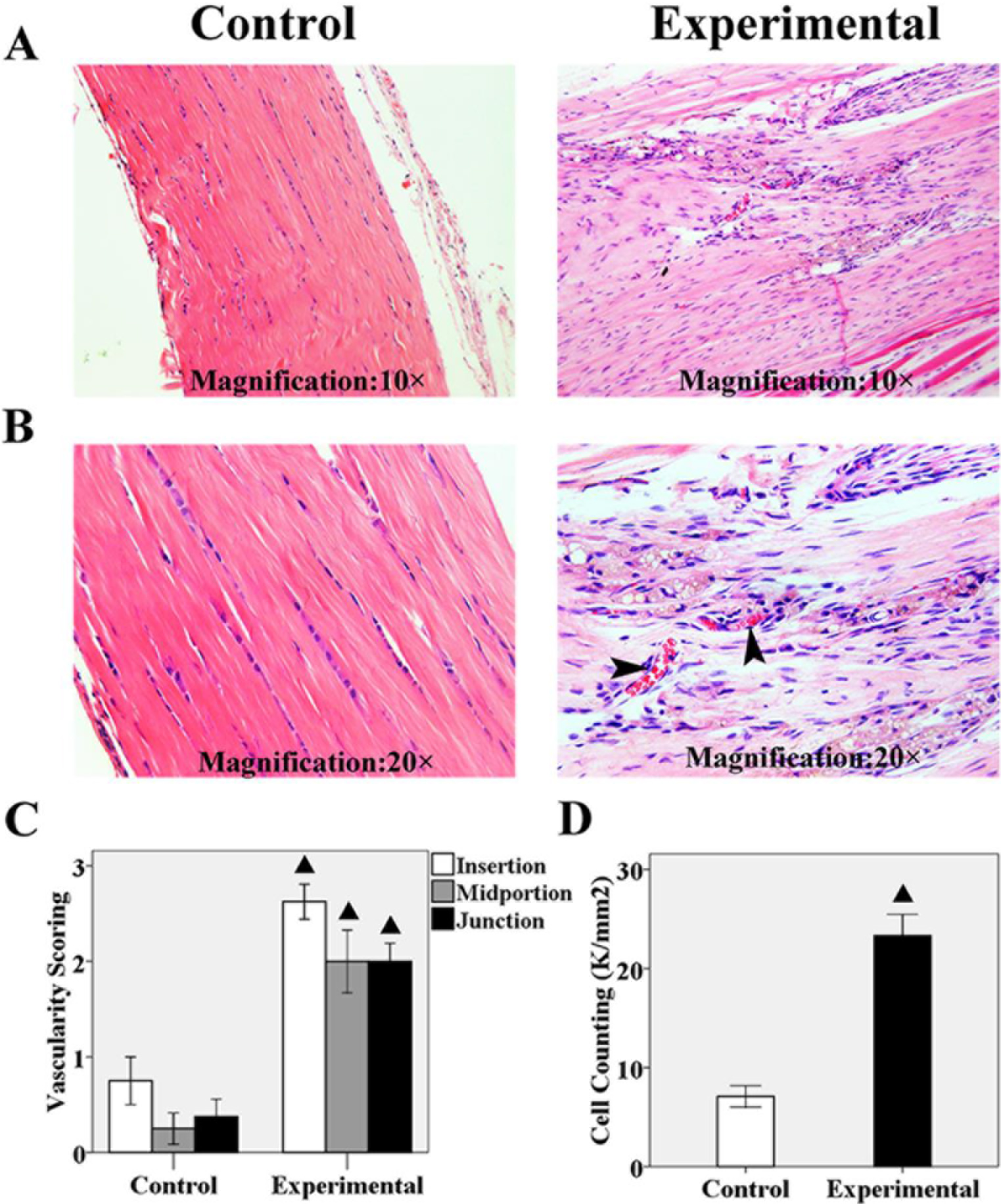
Hypercelluarity, neovascularization in the contralateral Achilles tendon after 12 weeks unilateral tenotomy. **A and B,** Representative H&E staining of Achilles tendon in experimental and control group. The hypercelluarity, fiber disorganization and neovascularization (shown with black arrows) was detected in the experimental group compared to control group. **C,** The vascularity scoring of experimental group in three portion of Achilles tendon was significant increased compared to the control group. **D,** The Achilles tenocytes counting of experimental group was significant more than control group. ▲ represent the comparison with control group, **P** < 0.05.

### Fiber disarrangement and collagen fusion in the contrallateral Achilles tendon after 12 weeks unilateral tenotomy

To comprehensively assess the microstructure of contralateral Achilles tendinopathy, we next performed both the SEM and TEM observation. The longitudinal collagen fiber disorganization was found in contralateral Achilles tendon versus control. There were more fused collagen fibers in the contralateral Achilles tendon with higher fiber diameter. Moreover, micro-gaps between fiber collagens were commonly observed (Fig.2). Interestingly, the ossification lesions with chondrocyte-like cells and neovasculars were detected inside the contralateral Achilles tendon compared to control (Fig.3), which could further contribute to the fiber collagen fusion and disorientation. These results confirmed the contralateral Achilles tendinopathy after 12 weeks unilateral tenotomy.

**Figure 2.**
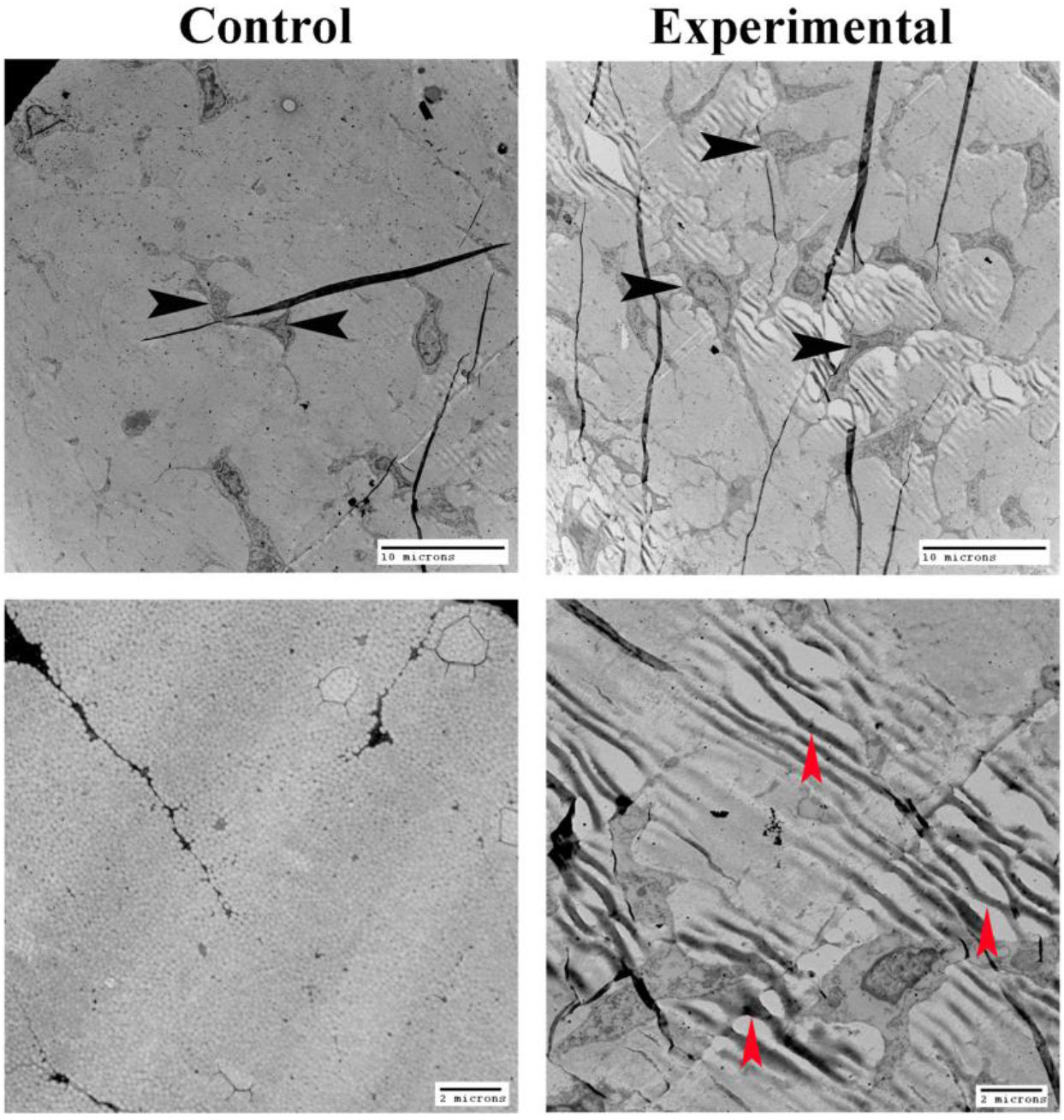
Representative Achilles tendon transmission electron microscopy observation. The longitudinal collagen fiber disorganization was found in ontralateral Achilles tendon versus control. There are micro-gaps between collagen fibers in the experimental group. Fiberswith higher diameters (shown with red arrows) were presented in the experimental group compared to control group. More tenocytes (shown with black arrows) were located in the experimental group.

**Figure 3.**
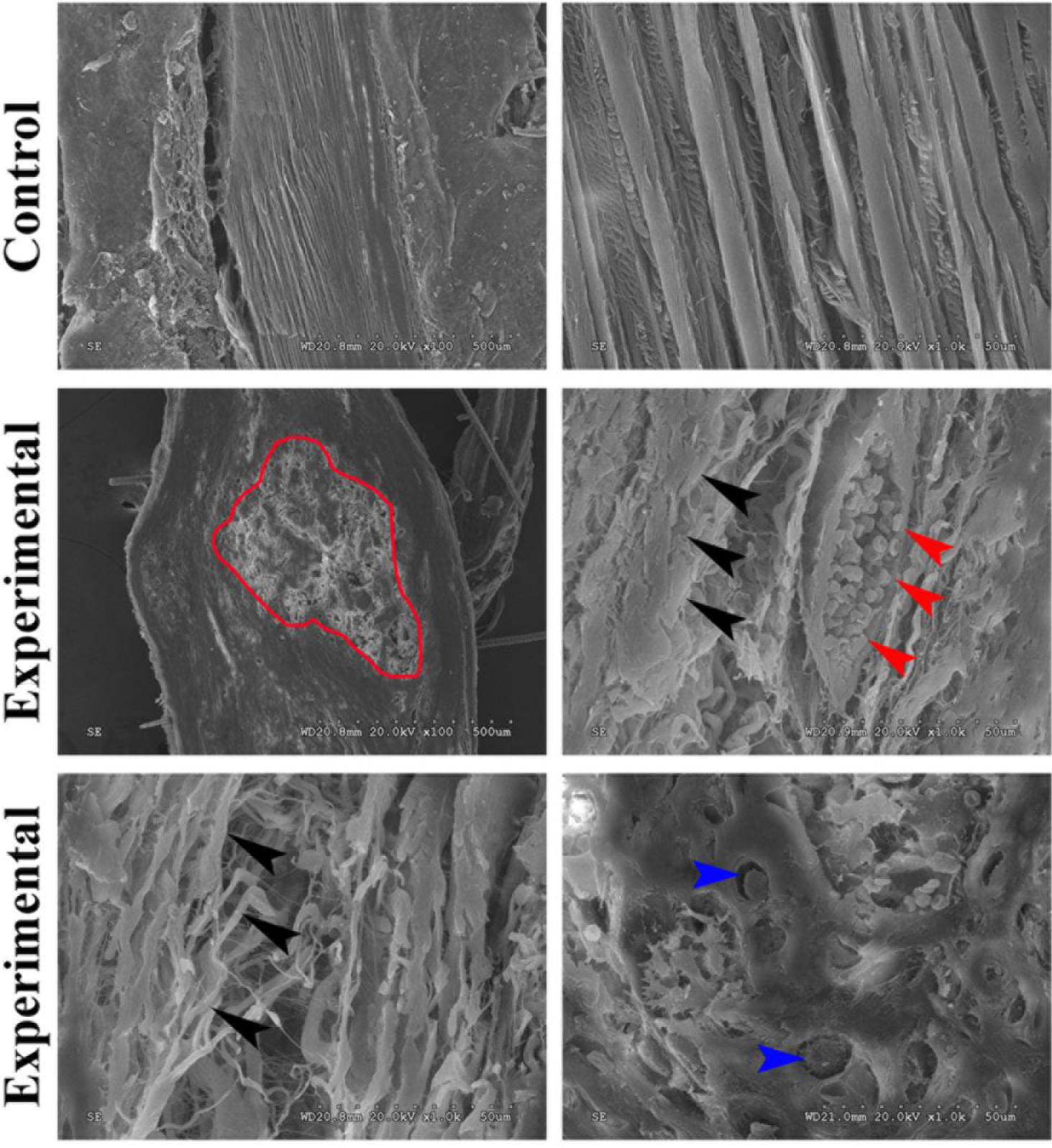
Representative Achilles tendon scanning electron microscopy observation. A ectopic ossification lesion (delineated by red line) was shown in the experimental Achilles tendon, chondrocytes-like cells with regular lacuna (shown with blue arrows) has been detected in the ossification area. New vascular growth (shown with red arrows) into Achilles tendon and longitudinal disoriented collagen fibers (shown with black arrows) was also demonstrated in the experimental group.

### Mineralization in the contralateral Achilles tendon and calcaneus deformity after 12 weeks unilateral tenotomy

To further evaluate the mineralization in the contralateral Achilles tendon, we examined and reconstructed the contralateral hindlimb using X-ray and micro-CT scanning. The ossifications were apparently detected in the both ends of contralateral Achilles tendon, which were mainly cortex bones with bone marrow cavity, whilst no ossification was detected in the control group. What’s more, in the experimental group, calcaneus deformity was presented in the reconstructed 3-dimension images. There were bony callus surround the calcaneus insertion of contralateral Achilles tendon. On the contrary, no calcaneus malformation was shown in the control group(Fig.4). These results strongly suggested mineralization was developed in the contralateral Achilles tendon, which is the most important feature of calcific Achilles tendinopathy.

**Figure 4.**
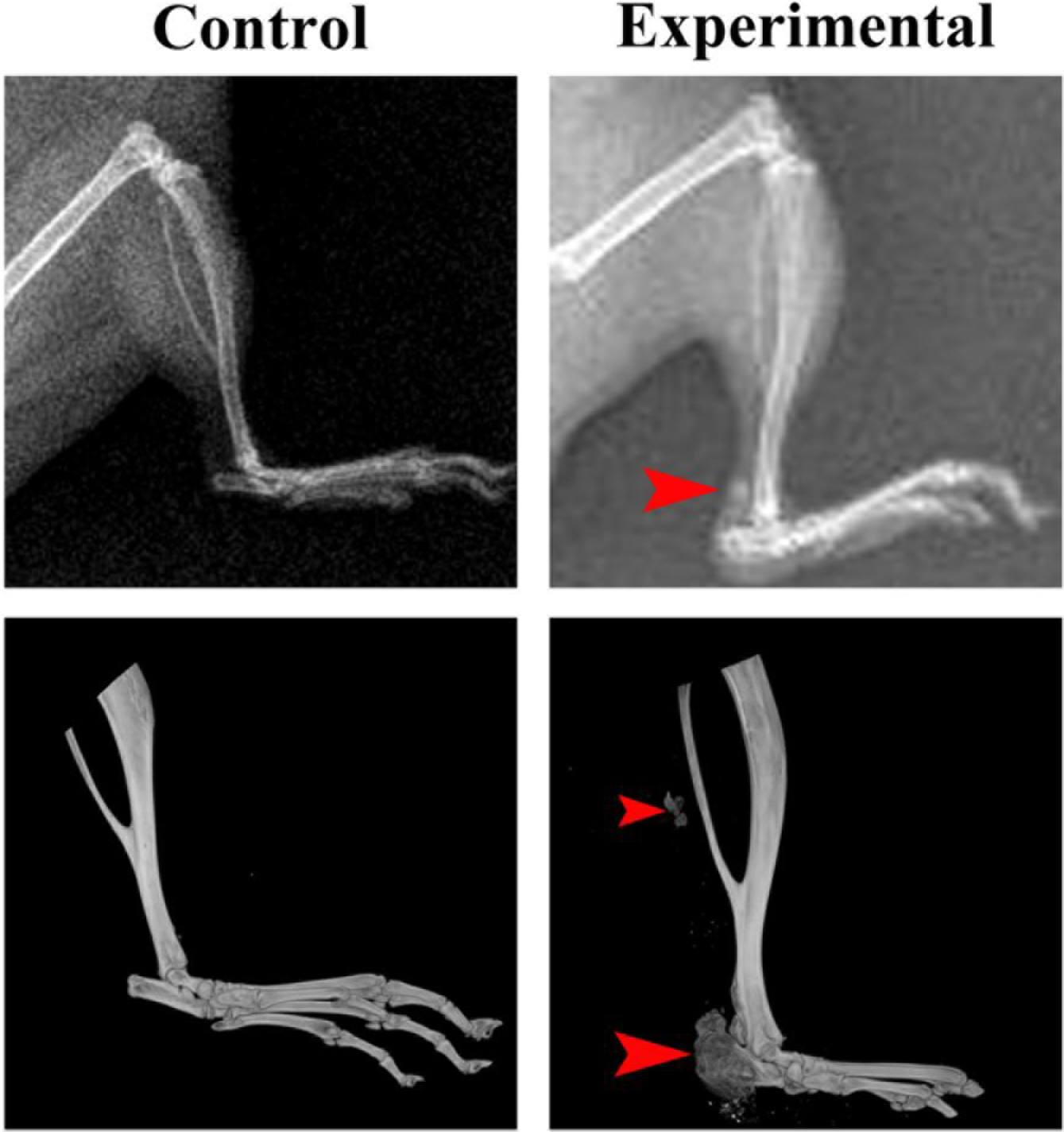
Representative observation of X-ray examination and micro-CT scanning for Achilles tendon of experimental group. A high density shadow of Achilles tendon in the experimental group has been detected by X-ray examination (shown with red arrows). Normal calcaneus shape was obscure from the X-ray plain film. Calcaneus deformity and Achilles tendon mineralization have been presented in the experimental group compared to control group by micro-CT three dimensional reconstruction

### Fibrocartilage metaplasia and endochondral ossification in the contralateral Achilles tendon after 12 weeks unilateral tenotomy

To investigate the underlying mechanism of ossification in the contralateral Achilles tendon, we performed H&E and toluidine blue staining. The results showed a large number of chondrocyte-like cells were presented inside the ossification area of Achilles tendon and calcaneus (Fig.5), which implied endochondral ossification may be involved in this process. Consequently, collagen II, collagen X IHC staining were conducted to identify endochondral ossification, both of which play a crucial role in endochondral ossification. There were abundant collagen II and collagen X-positive cells inside both the ossification area of Achilles tendon and calcaneus, conversely rare positive cells were determined in the control group (Fig.6). In addition, IF staining confirmed that more collagen III expression in the tenocytes of experimental group versus control, which indicated the achilles tendinosis in the experimental group(Fig.6). These results suggested that ossification of the contralateral Achilles tendon is cell-mediated endochondral ossification. Fibrocartilage metaplasia from either tenocyte stem cells or bone marrow stem cells could be the initial step of endochondral ossification in Achilles tendon enthesis.

**Figure 5.**
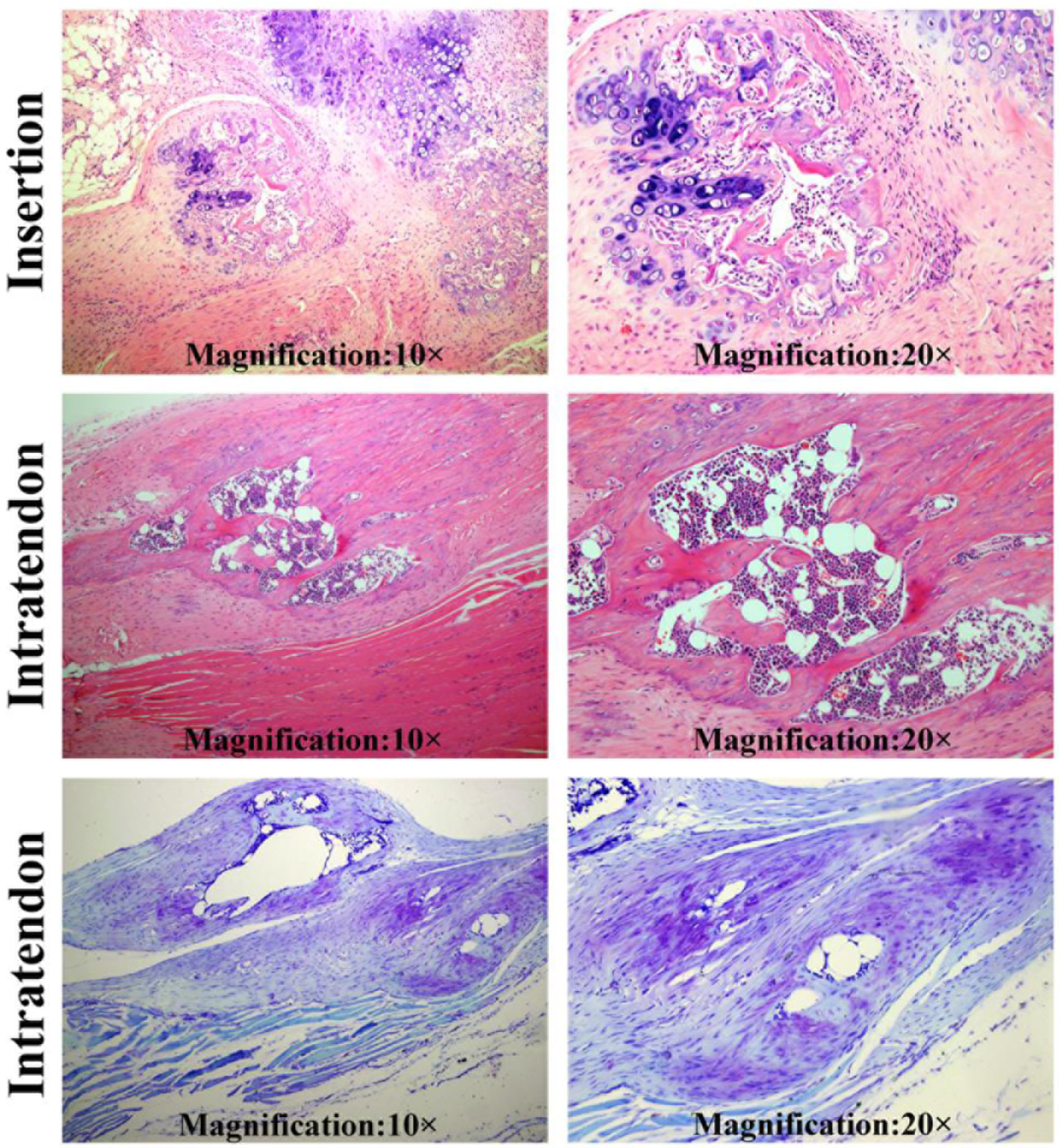
Fibrocartilage metaplasia and endochondral ossification in the contralateral Achilles tendon after 12 weeks unilateral tenotomy. Fibrocartilage metaplasia in the calcaneus insertion of experimental group has been found. Hypertrophic chondrocyte-like cells were distributed in the calcaneus and Achilles tendon in the experimental group. A mature cortex bone with bone marrow cavity has been detected intra the Achilles tendon. Ossification with specific stained chondrocytes intra Achilles tendon has been shown by toluidine-blue staining in experimental group.

**Figure 6.**
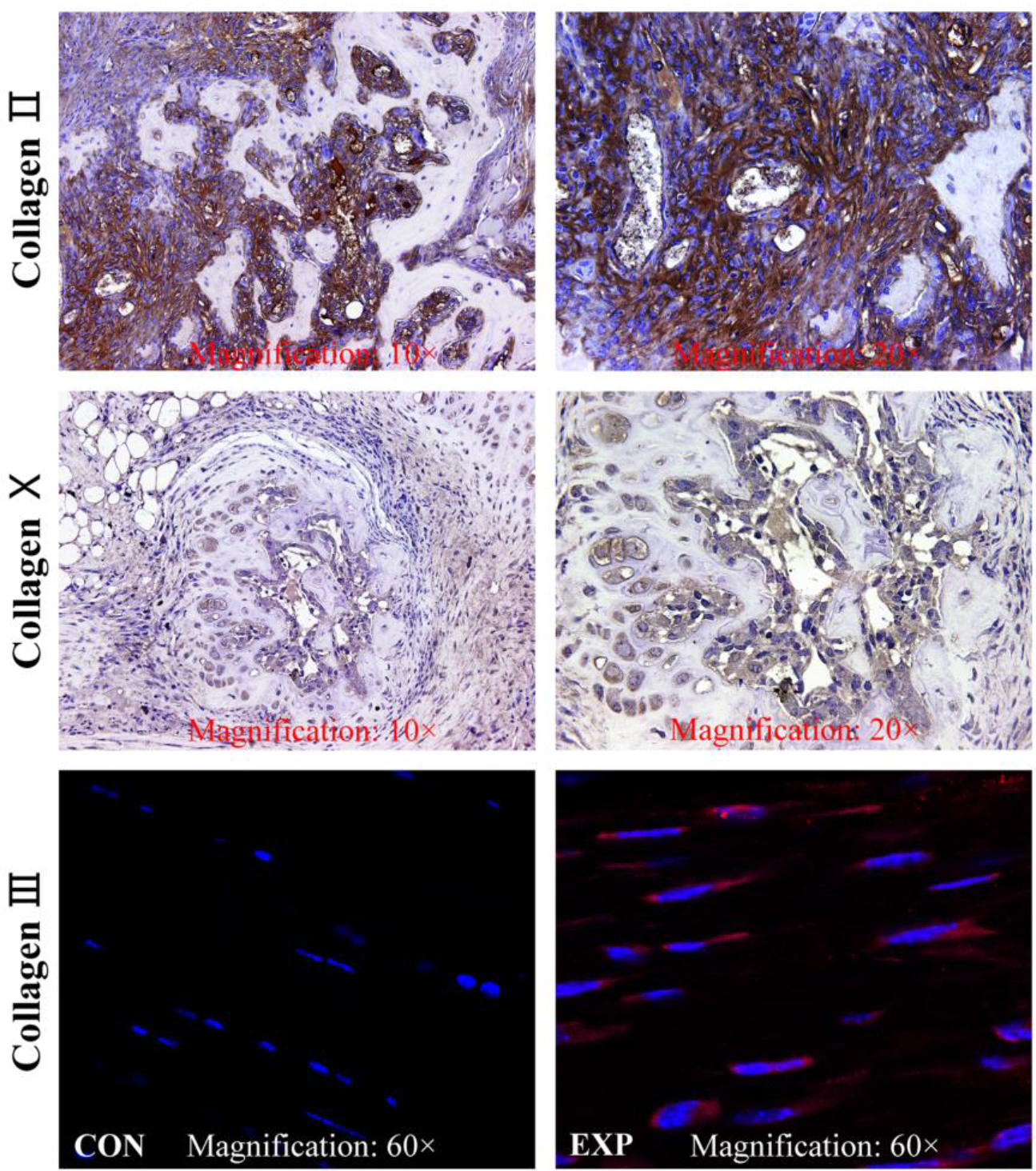
Representative IHC and IF staining for collagen protein expression in the experimental group. Collagen II and collagen X protein has been expressed in the ossification area. Large number of positive-stained cells presented in the ossification area of Achilles tendon and calcaneus. Collagen III expression by tenocytes in experimental group was detected in the Non ossification area of Achilles tendon compared to control group.

### Poor biomechanical properties in contralateral Achilles tendon after 12 weeks unilateral tenotomy

In general, Achilles tendinopathy results in poor biomechanical properties, which is considered the main reason for Achilles tendon rupture under normal loading[16]. We achieved the biomechanical analysis of contralateral Achilles tendon and recorded the maximal failure tensile loading and the stiffness of contralateral Achilles tendon. The cross-sectional area was decreased slightly in the experimental group. The maximal failure tensile loading in the experimental group was significantly reduced while the stiffness was significantly increased in the experimental group compared to control group(Table 1). The biomechanical properties changes may result from the ossification development and collagen fiber disorganization in the Achilles tendon. In summary, these results delineated the significant biomechanical properties loss in the contralateral Achilles tendon after 12 weeks unilateral tenotomy, which is the essential feature of Achilles tendinopathy.

**Table 1:** Biomechanical parameters of Achilles tendon complexFigure.

| Group | Control(n=8) | Experimental(n=8) | p Value |
| --- | --- | --- | --- |
| Cross-sectional area (mm<sup>2</sup>) | 0.29 ± 0.03 | 0.32 ± 0.04 | 0.08 |
| Failure tensile load (N) | 13.02 ± 0.86 | 8.35 ± 0.58 | 0.00 |
| Stiffness (N/mm) | 25.79 ± 2.01 | 32.00 ± 2.52 | 0.00 |

### Discussion

Calcific Achilles tendinopathy is a common and significant disease in elite athletes and older, sedentary and overweight individuals[17]. The aetiopathogenesis and treatment of this disease remains controversial[18], which gives rise to more interests in developing better animal models to address the gap in our understanding of calcific Achilles tendinopathy. Although unilateral Achilles tenotomy has been widely employed as a trauma-induced heterotopic ossification model in the previous studies[19–21], the changes of contralateral Achilles tendon in this model has not been reported. In the present study, we found the spontaneous contralateral Achilles tendinopathy after unilateral transection of Achilles tendon in mice, which could be a novel calcific Achilles tendinopathy model.

Compared to the existing animal models for Achilles tendinopathy investigation, our model has several benefits: firstly, the model of contralateral calcific Achilles tendinopathy is quite simple but highly reproducible which can be created only with the unilateral Achilles tendon transection. Non-special instrument is required, while some particular machines such as running-mills and pneumatic pistons[5,8,11,12], are generally necessary in the existing models. Secondly, it will be beneficial for imitating the natural pathogenesis of calcific tendinopathy in the contralateral Achilles tendons without any additional intervention. Furthermore, since this model has been established in mice, it is possible to take advantage of existing transgenic mice to investigate the pathogenesis of Achilles tendinopathy. With this model, the genetic predispostion of Achilles tendinopathy could be reconsidered instead of poorly understood.

Understanding the main pathophysiology underlying Achilles tendinopathy is the core to develop mechanism-based animal models[22]. Currently, there is still a long way to go before we comprehensively delineate the mechanism of calcific Achilles tendinopathy, however, overuse has generally been supposed as a critical extrinsic factor of Achilles tendinopathy[23]. Thus, several overuse-induced animal models of tendinopathy have been established [5,8,24]. In the present study, unilateral Achilles tendon transection leads to dysfunction of the injury tendon, subsequently, the equilibration of mechanical loading has been broke on contralateral Achilles tendon, where more stress loading are bearing while standing. To some extent, the contralateral Achilles tendon will encounter overuse. The overload stress induces the micro-injury of tendon to accelerate the degeneration of contralateral Achilles tendon. With time going by, unbalanced homeostasis in contralateral tendon will aggravate the Achilles tendinopathy subsequently. These results also strongly suggest that the contralateral Achilles tendon could not be indentified as the valid control of the unilateral tenotomy model in mice.

Interestingly, the morphological character of calcaneus has been remodeled in this model. From the results of Micro-CT analysis, the deformity of calcaneus has been detected, especially in the insertional region of Achilles tendon. The bony callus mass emerges in the posterosuperior area of calcaneus. The histological assessment indentify the fibrocartilaginous callus, woven bone and laminar bone with bone marrow cavity are located in the interface of the Achilles tendon enthesis. These results indicated the endochondral ossification is involved in the process of calcific Achilles tendinopathy, which is consistent with the study conducted by Lin et al in rats[25]. Consequently, we proposed calcaneus stress fracture could play an important role via endochondral ossification during calcific Achilles tendinopathy. The mechanical overload on the contralateral hindlimb induces stress fracture of calcaneus and increasing tensile loading of Achilles tendon results in calcaneus avulsion fracture where micro-fracture and micro-repair may occur coincide. Therefore, the calcaneus has been remodeling through endochondral ossification during the fracture healing.

The secondary blood vessel invasion into the tendon and potential fibrocartilage metaplasia form the tendon fibroblast may further contribute to the calcific deposition or ossification in the Achilles tendon. From this model, it is notable that calcaneus and Achilles tendon should be deemed to be a functional unit. The calcaneus fracture especially in the insertion of Achilles tendon may trigger subsequent calcific Achilles tendinopathy, which indicated the prevention of insertional Achilles tendinopathy should be taken into consideration during the treatment of calcaneus avulsion fracture or stress fracture treatment. Active precaution for calcaneus stress fracture may confer benefit to Achilles tendinopathy prevention.

Additionally, a solitary ectopic ossification is detected in the enthesis of soleus muscle from micro-CT analysis, indicating the erroneous chondral differentiation of tendon derived stem cells in the Achilles tendon may be involved. It has been proposed the formation of calcific deposits in calcifying tendinopathy is chondrocyte-mediated[26], which has also been demonstrated in histological analysis of this study. With mechanical loading overuse, the erroneous differentiation of TDSCs to chondrocyte and osteoblast could compromise the normal tendon healing, leading to weaken properties and activity-related pain[27]. Hence, the redirection of TDSCs differentiation to tendoncyte has been supposed to be a novel treatment target for Achilles tendinopathy[28]. Nevertheless, the underlying pathogenesis of aberrant TDSCs differentiation is not fully understood at the moment, which may restrict development of effective or evidence-based therapy strategy. Therefore, through by the model, it will facilitate our better understanding on the pathogenesis of TDSCs erroneous differentiation in the calcific Achilles tendinopathy, consequently encouraging a novel perspective for its treatment.

However, there are some drawbacks to this model indeed. Lack of internal control may result in overlook of the individual variance. It is important that the animal model should provide reproducible results that consistent with human conditions. When it refers to the Achilles tendinopathy in sedentary population, which means disuse or underuse of Achilles tendon, this model is not suitable. In addition, taking account of the crosstalk within the bilateral Achilles tendon and the existing evidence from bench and bedside, further studies should look into either the potential effect of sympathetic nerve modulation or endocrine predisposing factors in this model.

In conclusion, we hereby introduce a novel simple but reproducible spontaneous contralateral calcific Achilles tendinopathy model in mice, which represents the overuse conditions during the tendinopathy development in human-beings. It should be a useful tool to further study the underlying pathogenesis of calcific Achilles tendinopathy, including endochondral ossification from cancalneus stress fracture and the erroneous differentiation of TDSCs.

## Acknowledgements

This research was supported by grants from National Natural Sciences Foundation of China (31271271, 31228007, 81260401 and 81171764). We also thank Lu Tang and Aihong Yang for excellent technical assistance.

## Author contributions

All authors were involved in drafting the article or revising it critically for important intellectual content, and all authors approved the final version to be published. Dr. Dadi Jin and Dr. Xiaochun Bai had full access to all of the data in the study and takes responsibility for the integrity of the data and the accuracy of the data analysis. Study conception and design. Dadi Jin, Xiaochun Bai, Liang Wang, Minjun Huang. Acquisition of data. Liang Wang, Minjun Huang, Zhongmin Zhang, Tianyu Chen, Jian Jin, Xiaochen Zheng, Bin Huang, Bo Yan, Yuhui Chen, Shengfa Li. Analysis and interpretation of data. Liang Wang, Minjun Huang, Pinglin Lai. Manuscript preparation. Liang Wang, Minjun Huang, Dadi Jin, Xiaochun Bai.

